# ATP hydrolysis coordinates the activities of two motors in a dimeric chromatin remodeling enzyme

**DOI:** 10.1101/2020.11.17.387662

**Authors:** Stephanie L. Johnson, Geeta Narlikar

**Affiliations:** Department of Biochemistry and Biophysics, University of California, San Francisco, San Francisco, CA, USA

## Abstract

ATP-dependent chromatin remodelers are essential enzymes that restructure eukaryotic genomes to enable all DNA-based processes. The diversity and complexity of these processes are matched by the complexity of the enzymes that carry them out, making remodelers a challenging class of molecular motors to study by conventional methods. Here we use a single molecule biophysical assay to overcome some of these challenges, enabling a detailed mechanistic dissection of a paradigmatic remodeler reaction, that of sliding a nucleosome towards the longer DNA linker. We focus on how two motors of a dimeric remodeler coordinate to accomplish such directional sliding. We find that ATP hydrolysis by both motors promotes coordination, suggesting a role for ATP in resolving the competition for directional commitment. Furthermore, we show an artificially constitutive dimer is no more or less coordinated, but is more processive, suggesting a cell could modulate a remodeler’s oligomeric state to modulate local chromatin dynamics.

## 1 Introduction

ATP-dependent chromatin remodelers are enzymes that restructure eukaryotic genomes in order to facilitate all DNA-based processes, including transcription, DNA damage repair, and replication ([1, 2, 3]). Their substrates are chromatin, the protein-nucleic acid complex that stores a cell’s genetic information, and in particular the nucleosome, the basic repeating unit of chromatin, consisting of 147 bp of DNA wrapped nearly twice around a core of histone proteins. Chromatin remodelers catalyze a variety of transformations of nucleosomes, including nucleosome sliding, nucleosome assembly and disassembly, and the exchange of core histone proteins with histone variants ([4]).

Perhaps not surprisingly, given the involvement of remodelers in so many key genomic processes, mutations or disruptions to remodelers have been implicated in cancer, developmental disorders, and other diseases ([1, 5, 6, 7]). Yet they remain elusive therapeutic targets because they are challenging to study by the conventional biochemical approaches that have rendered other molecular complexes amenable to pharmaceutical analysis. In addition to the inherent complexity of their chromatin substrates, remodelers themselves can be large, multi-subunit complexes, up to more than a megadalton in size, and their component subunits can have multiple “moving parts” that undergo large conformational changes during the reaction cycle ([4, 8, 9]). Moreover, the reactions these remodelers catalyze often involve large-scale physical, not just chemical, alterations to the substrate ([4]).

In light of these challenges, single molecule biophysical approaches have generated significant enthusiasm in the field for their potential to open new windows into remodelers ([10, 11, 12, 13, 14, 15, 16, 17, 18]). These approaches overcome some of the difficulties listed above by bypassing the need to study remodelers in asynchronous ensemble populations. In addition, many single molecule biophysical techniques are more readily suited to detecting large physical or structural changes to a substrate.

Here we use single molecule fluorescence resonance energy transfer (smFRET) to gain detailed mechanistic insights into a common but still opaque reaction catalyzed by many chromatin remodelers: directional nucleosome sliding. A majority of chromatin remodelers studied to date are capable of sliding nucleosomes along the DNA, and, interestingly, most of them do so in a highly regulated manner, preferentially sliding a nucleosome towards the longer flanking DNA ([4, 19]). This directional nucleosome sliding is thought to contribute to the generation of evenly spaced nucleosomal arrays, as nucleosomes are continuously moved in the direction of the longer linker DNA until the lengths of DNA between nucleosomes has been equalized ([20, 21]). Evenly spaced arrays of nucleosomes are associated with transcriptionally silenced, heterochromatic regions of the genome, as well as with other chromatin structural features like TAD boundaries ([22, 23, 24, 25, 26]). Nucleosome sliding in the direction of longer flanking DNA may also allow remodelers to facilitate transcriptional activation and/or repression, by moving nucleosomes away from genomic features such as DNA-bound transcription factors ([27, 28]).

The ISWI family of chromatin remodelers has become a paradigm for this directional nucleosome sliding activity, particularly the ACF remodeler, a complex of SNF2h, the ATP-hydrolyzing motor subunit, and Acf1, a non-catalytic accessory subunit. A common in vitro proxy assay for the activity of evenly spacing nucleosome arrays is the centering of a mononucleosome, a single nucleosome on a short DNA: equalizing the linker DNAs in an array translates to equalizing the lengths of the DNAs flanking the mononucleosome. In ensemble in vitro experiments using this mononucleosome proxy, both ACF and the SNF2h motor subunit alone slide mononucleosomes faster when they have longer flanking DNA ([29, 21]). This kinetic discrimination explains their preference for sliding nucleosomes in the direction of the longer flanking DNA, and, presumably, their ability to space arrays ([21]).

In addition to its biological importance, the ability to slide nucleosomes preferentially towards longer flanking DNA presents a fascinating biophysical challenge. Not only must the length of the DNA external to each nucleosome be assessed by these enzymes, but a *comparative* measurement of the *relative* lengths of DNA on either side of the nucleosome must be accomplished. How is this comparative assessment made? Part of the answer seems to lie in the ability of ACF and SNF2h, though monomeric in solution, to dimerize on the nucleosome, and in fact to slide nucleosomes most efficiently as dimers ([30, 11, 31]). However, this raises the further question of how two motors coordinate their activities across their substrate without engaging in a “tug-of-war”.

To better understand how this coordination is achieved, we capitalized on our recent work with synthetic, constitutively dimeric forms of the SNF2h motor subunit ([31]). These synthetic enzymes, which we call [wt]-[wt], are covalently linked such that they are dimeric in solution, which allows us to also make asymmetric mutations in the [wt]-[wt] complex, to investigate various aspects of the two protomers’ coordination.

These synthetic dimers, in conjunction with the ability to watch them individually remodel single nucleosomes by smFRET, enabled us to dissect how two SNF2h motors coordinate their activities in unparalled detail. We show that covalently linking two SNF2h motors not only maintains their ability to coordinate their activities, but also makes the synthetic dimer processive, whereas SNF2h alone is not processive. Further, we show that ATP hydrolysis by both motors in a SNF2h dimer enables them to coordinate their DNA length sensing activities and avoid a tug of war. The use of ATP hydrolysis to regulate remodeling, not just as a source of energy coupled to the physical work of nucleosome sliding, is emerging as a common theme in DNA-length-sensitive nucleosome sliding enzymes ([15, 11, 12, 31]. We describe here new mechanistic details for how ATP can be used to regulate nucleosome sliding in a paradigmatic remodeler family.

## 2 Results

### 2.1 A constitutively dimeric SNF2h remodels single nucleosomes like wild-type SNF2h

Our previous work with the constitutively dimeric [wt]-[wt] construct uncovered no major remodeling defects compared to wild-type SNF2h in ensemble assays ([31]). However, single-molecule FRET provides a more detailed view of the nucleosome sliding reaction and can report on dynamics that might be obscured by the population averaging of ensemble assays. Therefore, we first compared the remodeling reaction of [wt]-[wt] to SNF2h at the single nucleosome level. We find that the [wt]-[wt] construct behaves similarly to SNF2h, with a key exception that will be discussed in the next section.

ISWI-family remodelers, including SNF2h, have been shown to slide single nucleosomes in an alternating pattern of pause and nucleosome translocation events ([11, 12, 13, 9]). When observed by single molecule FRET (Fig. 1(A)), the pauses appear as relatively long periods where the FRET remains constant and the nucleosome is not being slid (p_wait_, p_1_, and p_2_ in Fig. 1(B)), whereas nucleosome translocation appears as short, rapid drops in FRET intensity (called t_1_ and t_2_). Importantly, the pauses have been shown to be regulatory events. It is during the pause phases of the reaction that regulatory information such as flanking DNA length is sensed ([13, 9]) and, presumably, used to gate the sliding reaction that takes place in the translocation phases. In particular, the “decision” about which direction to move the nucleosome, based on regulatory information such as the relative lengths of the DNAs flanking the nucleosome, must happen in the pause, before the nucleosome is moved in the translocation phase.

**Figure 1:**
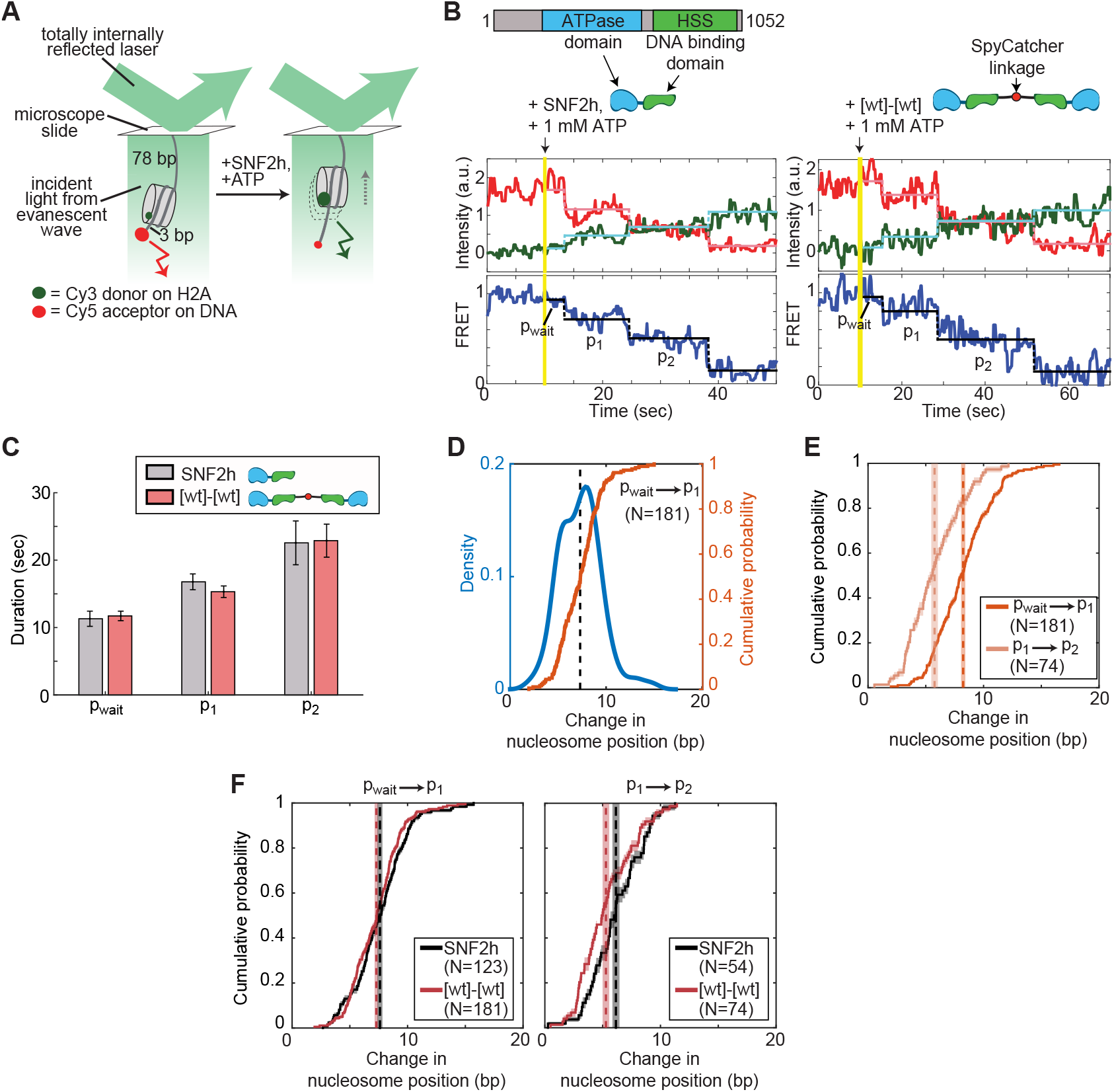
A constitutively dimeric SNF2h retains key features of the remodeling reaction at the single nucleosome level. (A) Schematic of the smFRET setup. Nucleosomes (here, initially end-positioned “3/78”) start in high FRET. As SNF2h slides the nucleosome towards the center of the DNA, the distance between the Cy3 and Cy5 dye pair increases, leading to decreased FRET. (B) (top) Domain architecture of SNF2h, showing the two major domains that will be drawn schematically in blue (ATPase domain) and green (DNA binding domain) in the rest of this work. (bottom) Example traces of SNF2h (left) and [wt]-[wt] (right) remodeling single nucleosomes, with the first three pauses labeled (p_wait_, p_1_, p_2_). Vertical yellow lines indicate time at which enzyme and ATP are injected into the sample chamber. (C) Average durations of the first three pauses exhibited by SNF2h or [wt]-[wt], for ∼100 nucleosomes each. Errors are bootstrapped as described in the Methods. (D) Kernel density estimation (KDE, blue) and empirical cumulative distribution function (CDF, orange) of the change in nucleosome position between p_wait_ and p_1_ for 181 nucleosomes remodeled by [wt]-[wt]. KDEs can be more intuitive visually, but CDFs are a more quantitative way to compare data sets. Peaks in the KDE correspond to steep slopes in the CDF. Vertical black dashed line is the mean. N: number of events included in the KDE and CDF. (E) CDFs of the change in nucleosome position between pwait and pi (first translocation phase) versus p_1_and p_2_ (second translocation phase) for [wt]-[wt], showing the initial larger step followed by a smaller second step. Mean step sizes are indicated by dashed vertical lines (with shaded region representing the error) and are 7.3±0.2 and 5.3±0.3 bp, for the first and second translocation phases respectively. (F) CDFs of the change in nucleosome position during the first (left) or second (right) translocation phases for SNF2h versus [wt]-[wt]. Mean step sizes for SNF2h are 7.6±0.2 and 6.2±0.3 bp for the first and second translocation phases respectively. See Fig. S5 for corresponding step size KDEs. In all panels, enzyme concentrations are saturating (51 nM SNF2h, 25 nM [wt]-[wt]); ATP concentration is also saturating (1 mM). Errors on the CDFs and mean step sizes are determined by a bootstrapping approach (see Methods).

This also means that any *coordination* between the two SNF2h motors to “decide” which direction to slide the nucleosome must take place during the pauses. As shown in Fig. 1(C), wild-type SNF2h and the constitutive [wt]-[wt] dimer have identical pause durations when they remodel initially end-positioned “3/78” nucleosomes (nucleosomes with 3 bp of flanking DNA on one side and 78 bp on the other). This indicates that by forcing SNF2h to be a dimer, we have not compromised the ability of the two protomers to coordinate their activities; a compromise in coordination should result in a “tug-of-war” that would make it harder for the enzyme to exit the pause phase, and thus should increase the durations of the pauses. We do not observe any such increase in pause duration with [wt]-[wt]. (The pause durations are not shorter with [wt]-[wt], either, indicating that we have not created a more efficient remodeler.)

In addition to a stereotyped, alternating pattern of pauses and translocations, all ISWI-family remodelers studied by smFRET to date also exhibit a stereotyped pattern of step sizes, defined as the distance the nucleosome is slid during the translocation phases. The first translocation event slides the nucleosome ~7-8 bp on average, whereas subsequent translocation events each move the nucleosome ~5 bp (Fig. 1(D,E), Fig. S5, [11, 12, 13, 9]). This pattern of an initial large step followed by smaller steps is maintained in the constitutive dimer (Fig. 1(F)).

Thus by forcing SNF2h to be a constitutive dimer, we have not compromised the efficiency of the remodeler’s escape from the regulatory pause phase, nor its ability to properly slide the nucleosome in the translocation phases. There are a number of ways that two enzymes could have the same overall remodeling rate when measured at the ensemble level and yet could differ at the single nucleosome level—e.g., [wt]-[wt] could have exhibited longer pauses but also larger step sizes—but this is not what we observe. Instead, covalently linking the two HSS domains of a SNF2h dimer has no effect on the two motors’ abilities to efficiently slide single nucleosomes, and so the [wt]-[wt] construct can be used to probe protomer coordination.

### 2.2 Nucleosome sliding catalyzed by a constitutively dimeric SNF2h is more processive than by wild-type SNF2h

A major advantage that smFRET has over ensemble assays for measuring nucleosome sliding is that smFRET can measure the *processivity* of the remodeling enzyme. We define processivity as the number of pause-translocation-pause-translocation cycles that the enzyme can catalyze before dissociating from the nucleosome, or, equivalently, how long the enzyme can continue to slide a nucleosome under chase conditions. Previous single molecule work with the ISWI remodeler ACF, a complex of SNF2h and the accessory subunit ACF1, showed that ACF is highly processive ([11]). We asked whether the same is true of the motor subunit alone, and whether the enzyme’s processivity is affected in the constitutively dimeric construct.

To quantify processivity, we generated nucleosomes initially positioned in the center of a long DNA, with 60 bp on each side (“60/60” nucleosomes; Fig. 2(A)). Although 60 bp is longer than SNF2h’s length sensitivity of 25-30 bp ([21, 31, 32, 33]), by an ensemble gel remodeling assay, almost 40% of the population of nucleosomes is still near the center of the DNA after the ~3-5 minutes we can image nucleosomes before the FRET dyes photobleach (Fig. S3(A)). The long flanking DNA ensures that the nucleosome remains sufficiently far from the surface of the microscope slide to prevent potential artifacts from nucleosome-surface interactions.

**Figure 2:**
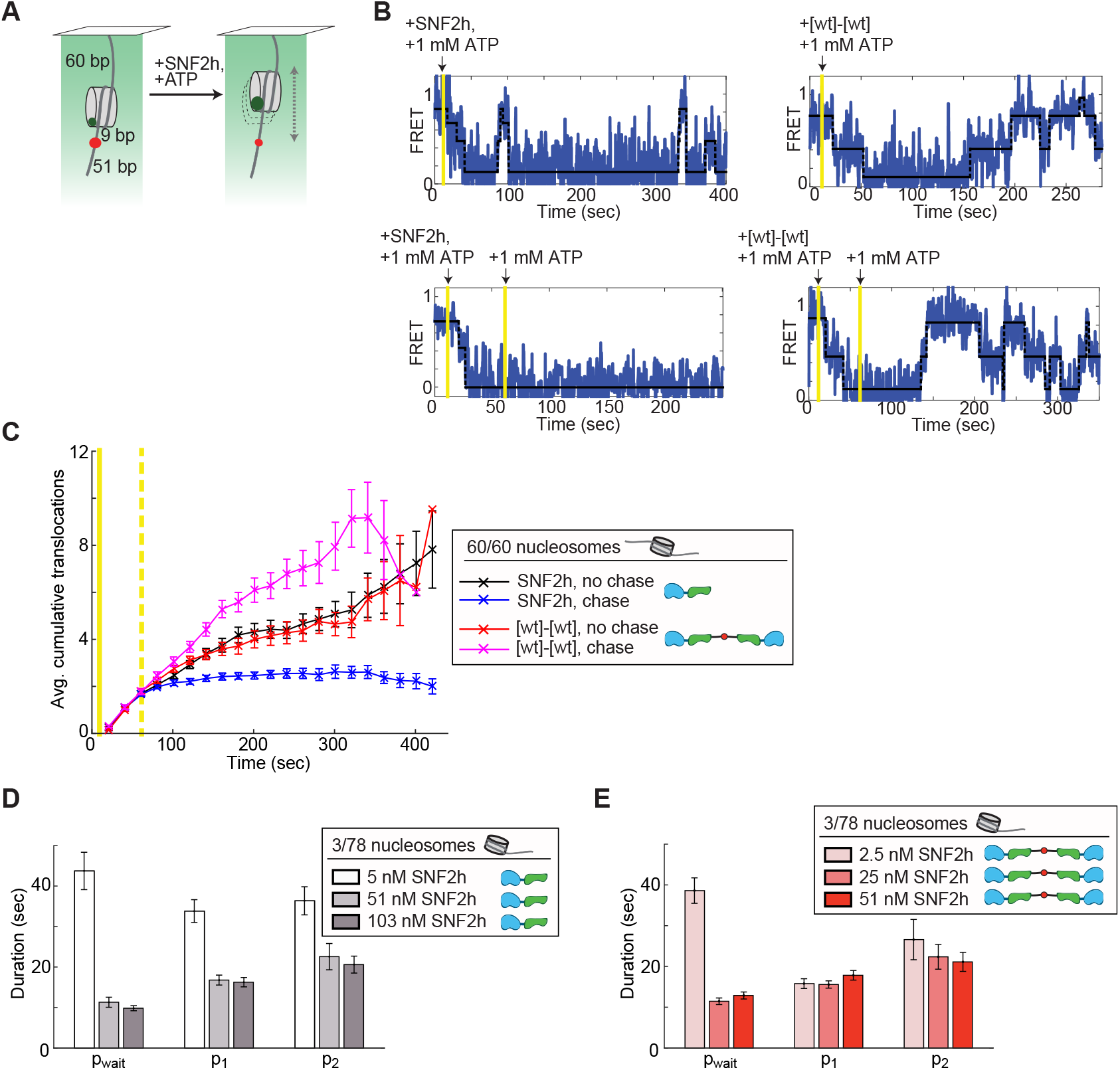
A constitutively dimeric SNF2h exhibits increased processivity compared to wild type SNF2h. (A) Schematic of the smFRET assay with initially centered 60/60 nucleosomes. (B) Example traces of SNF2h vs. [wt]-[wt] on 60/60 nucleosomes, without (top) or with (bottom) a 1 mM ATP chase about 1 minute after the first injection to start remodeling. [wt]-[wt] is significantly more likely to continue to remodel a nucleosome after a 1 mM ATP chase than is SNF2h, indicated by the repeated cycles of increasing and decreasing FRET in the bottom right trace. (C) Quantification of processivity of SNF2h vs. [wt]-[wt]. An accumulation of translocations over time indicates that remodeling continues. The error increases with time because fewer trajectories survive without photobleaching at longer time points. Solid vertical yellow line indicates injection of enzyme plus ATP; dashed vertical yellow line indicates the timepoint of the chase, if applicable. Enzyme concentrations here and in (B) were 103 nM SNF2h and 51 nM [wt]-[wt], with 1 mM ATP. See Fig. S4 for the cumulative translocations in individual traces that went into these averages. (D) Average durations of the first three pauses of the SNF2h remodeling reaction as a function of SNF2h concentration, for ∼100 nucleosomes at each concentration. At subsaturating SNF2h (5 nM), the durations of all three pauses are longer, indicating that binding or re-binding of SNF2h becomes rate-limiting. (E) Average durations of the first three pauses of the [wt]-[wt] remodeling reaction as a function of [wt]-[wt] concentration, for ∼100 nucleosomes at each concentration. Here only the first pause duration increases at the sub-saturating enzyme concentration, indicating that re-binding of [wt]-[wt] is not rate-limiting for the p_1_ and p_2_ pauses. ATP concentration here and in (D) was 1 mM. Errors were bootstrapped as described in the Methods.

We measured remodeling of these 60/60 nucleosomes by SNF2h and [wt]-[wt] under chase conditions, by injecting enzyme and ATP into the sample chamber, allowing remodeling to commence, and then flushing the chamber with an excess of buffer containing ATP but no additional enzyme (see Methods). Any enzyme that dissociates from its substrate will experience near-infinite dilution into the large volume of buffer above the surface-attached nucleosomes, and so any remodeling events after the second buffer exchange must be carried out by enzymes that remain bound to the nucleosome. A processive enzyme will remain bound and will continue to remodel the nucleosome after the chase.

As shown in Fig. 2(B), after the injection of saturating ATP and either saturating SNF2h or saturating [wt]-[wt] into the sample chamber (first yellow bar), the 60/60 nucleosomes are repeatedly slid back and forth along the DNA, moving in and out of FRET range. Remodeling was either allowed to proceed normally, or a chase was performed into 1 mM ATP alone, without enzyme (second yellow bar in bottom example trajectories), about 1 minute after the initial injection of remodeler and ATP to start the reaction.

Interestingly, we find that SNF2h alone, unlike the ACF complex, is not processive. As shown in Fig. 2(C), in the absence of the chase, SNF2h continues to remodel the 60/60 nucleosomes past the center of the DNA, and so translocation events continue to accumulate with time. However, under chase conditions, very few trajectories continue to accumulate new translocation events after the chase—that is, remodeling stops for most nucleosomes.

On the other hand, [wt]-[wt] *does* continue to remodel 60/60 nucleosomes even under chase conditions, as shown by the continued accumulation of transitions even under chase conditions (magenta curve in Fig. 2(C)). Thus by making a constitutively dimeric SNF2h, we have made it more processive, more like the ACF complex.

Given how low SNF2h’s processivity is, does SNF2h dissociate from the nucleosome frequently, perhaps even after every pause-translocation cycle? To ascertain whether this is the case, we returned to the 3/78 nucleosome construct, for which more of the reaction is in FRET range, and asked how many of the pauses are sensitive to SNF2h concentration. Previous work with ACF showed that only the first pause is sensitive to enzyme concentration, meaning only the first pause has an enzyme binding event, consistent with ACF being highly processive ([11]).

As shown in Fig. 2(D), at a sub-saturating SNF2h concentration, all pause durations are longer than at saturating SNF2h, indicating that enzyme (re-)binding is rate-limiting for all pauses when SNF2h is sub-saturating. For [wt]-[wt], on the other hand, only the first pause is sensitive to enzyme concentration (Fig. 2(E)). Thus the SNF2h reaction can include a dissociation and re-binding event at every pause, whereas [wt]-[wt], like ACF, is highly processive and can catalyze multiple pausetranslocation cycles without dissociating.

The possibility that SNF2h can dissociate from its substrate at every pause has implications for the step size. As discussed above, ISWI enzymes exhibit a stereotyped behavior in that the first translocation event moves the nucleosome about twice as far as the second translocation event. This means that SNF2h “remembers” that the first translocation event has been completed and the next translocation should be shorter. Is this memory retained at sub-saturating SNF2h, during which at least one SNF2h protomer can dissociate, and the complex must wait for another to re-bind?

As shown in Fig. S5, this memory is indeed retained even at sub-saturating concentrations of SNF2h. In particular, at sub-saturating concentrations of SNF2h, the second step does not get longer. If the memory of the first step were were affected, we would expect the second step to be longer, more like the first step, or at least for there to be a population of nucleosomes with two long first-translocation-like steps. Instead, at sub-saturating SNF2h, the second step is, if anything, shorter than under saturating conditions. We speculate that at 5 nM SNF2h, one protomer remains bound to the nucleosome, and that one protomer alone can retain the memory of how far the next translocation event should slide the nucleosome. This memory could be enforced through the octamer distortion that we and others have recently described for ISWI enzymes ([16, 34, 35]), which is present even with a SNF2h monomer bound to the nucleosome ([16]).

In summary, although the [wt]-[wt] construct shares most features of the single-nucleosome remodeling reaction with SNF2h, the constitutive dimer is more processive, more like ACF. This increased processivity will be essential for constraining our model of protomer coordination by SNF2h after considering several asymmetric mutants below, and has implications for the roles of accessory subunits in modulating SNF2h’s activity, which will be addressed in the Discussion.

### 2.3 ATP hydrolysis coordinates the length-sensing activities of the two SNF2h motors

A fascinating observation from earlier smFRET studies with ISWI remodelers is that ATP is required for *both* the pause phases *and* the translocation phases of the remodeling reaction ([11, 12]). During the translocation phases, the enzyme is doing physical work to slide the nucleosome, and so a requirement for ATP hyrolysis makes sense. However, the role of ATP in the pauses remains unclear.

Since, as noted above, it is in the pause phases that the two SNF2h motors must jointly “decide” which direction to translocate the nucleosome, we speculated that ATP hydrolysis might be involved in protomer coordination. Of course, making a SNF2h mutant that is compromised in its ability to hydrolyze ATP cannot provide insight into this question, because such a mutant would have no remodeling activity at all. But with the constitutively dimeric [wt]-[wt] construct, we can mutate the ATPase domain of only one of the two protomers, and ask what effect such an asymmetric mutation has on nucleosome sliding. We call this construct [wt]-[WB], because one protomer has a mutation in the Walker B (WB) domain of the motor’s active site that compromises ATP hydrolysis ([31]).

The simplest hypothesis for what we might observe with such an asymmetric mutant would be two populations of remodeled nucleosomes, depending on the initial binding orientation of the [wt]-[WB] construct. ~50% of a population of an end-positioned 3/78 nucleosomes should be bound by [wt]-[WB] such that the wild-type protomer can slide the nucleosome towards the longer flanking DNA. We would expect remodeling by this population to proceed normally, at least for the first several rounds of pauses and translocation events. The other half of the nucleosomes will be bound by [wt]-[WB] in an orientation in which the catalytically compromised protomer should be the one to slide the nucleosome. In the extreme case, this population would not remodel at all, at least until the enzyme dissociates and re-binds in the other orientation. As discussed in the previous section, dissociation by the constitutive dimer is quite slow. So at the single nucleosome level, we might observe a population of nucleosomes that is remodeled normally, and a population that remodels too slowly (if at all) to be observed by smFRET.

As shown in Fig. 3, this is not what we observe. The durations of the pauses with [wt]-[WB] are at least twice as long as with [wt]-[wt], while other aspects of the reaction are unaffected. This means that a catalytically compromised protomer has a dominant negative effect on the wild-type protomer, preventing efficient exit from the pauses, while leaving the actual process of nucleosome sliding unaffected. The slow dissociation rate of the constitutive dimer rules out a model in which pauses are longer with [wt]-[WB] due to a need to wait for the enzyme to dissociate and re-bind in a productive orientation during each pause. (However, the ability of the [wt]-[WB] construct to center a population of nucleosomes at the *ensemble* level in [31], albeit 5 times slower than [wt]-[wt], does require dissociation and re-binding such that the wild-type promoter can eventually slide all of the nucleosomes to the center of the DNAs. The processivity data in Fig. 2(C) are consistent with a dissociation rate on the timescale of the ensemble remodeling reactions of [31], since the rate of accumulation of translocations does decrease with time.)

**Figure 3:**
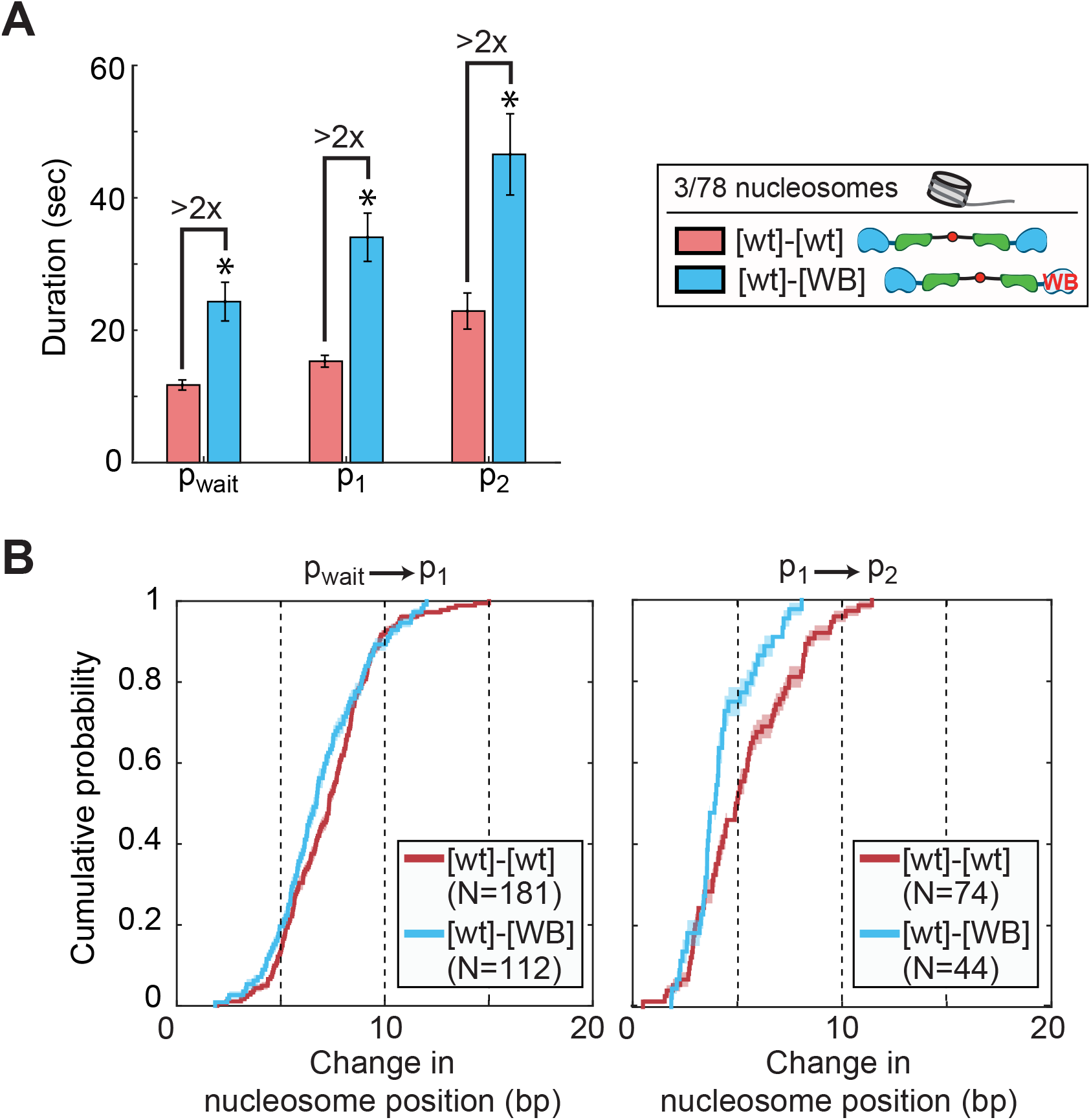
ATP hydrolysis is required for the two motors to coordinate and avoid a tug of war. (A) Average durations of the first three pauses when [wt]-[wt] or [wt]-[WB] remodels end positioned (3/78) nucleosomes, for ∼100 nucleosomes each. The asterisks indicate that these pause durations are lower bounds on the actual pause durations; the slow remodeling rate of [wt]-[WB] is masked by the competition with the rate of photobleaching (see Fig. S6 and [9]). (B) CDFs of the change in nucleosome position during the first (left) or second (right) translocation phases for [wt]-[wt] versus [wt]-[WB]. Mean step sizes for [wt]-[wt] are 7.3±0.2 and 5.3±0.3 bp for the first and second translocation phases respectively, and for [wt]-[WB] are 6.9±0.2 and 4.2±0.2 bp respectively. See Fig. S7 for corresponding step size KDEs. In all panels, enzyme concentrations are saturating (25 nM [wt]-[wt], 50 nM [wt]-[WB]); ATP concentration is also saturating (1 mM). Errors on the CDFs and mean step sizes are determined by a bootstrapping approach (see Methods).

In addition to pause durations, the step sizes with [wt]-[WB] differ slightly from those observed with [wt]-[wt]. Specifically, the second step is slightly shorter with [wt]-[WB] than with [wt]-[wt] (Fig. 3(B)). However, the overall behavior of an initial large step followed by a step that moves the nucleosome about half the distance of the first step is retained. This makes [wt]-[WB]’s step size different from the altered step sizes that we previously observed with a mutation to the acidic patch on the nucleosomal surface ([9]). Mutating the acidic patch, which also leads to longer pause durations, is so far the only mutation that breaks the 7-8 bp first step, 3-5 bp second step pattern observed with all ISWI enzymes. It remains unclear why this pattern of step sizes is so robust and so highly conserved in these enzymes (and possibly conserved within the broader superfamily of remodelers, e.g. CHD4 [36]).

Returning to the question of protomer coordination, the longer pause durations with [wt]-[WB] suggest that the WB mutation does indeed affect this coordination: a catalytically compromised protomer can prevent its wild-type partner from efficiently exiting the pause phase and sliding the nucleosome. Since the two protomers are making a coordinated “decision” about which direction to slide the nucleosome in response to the lengths of DNA flanking the nucleosome, we next introduced a mutation to abolish one protomer’s ability to sense flanking DNA. Specifically, we removed the HSS domain, the domain of SNF2h that binds to and senses flanking DNA (Fig. 1(B)), from the catalytically compromised protomer, to make a construct called [wt]-[WB/ΔHSS].

Surprisingly, the removal of the HSS from the catalytically compromised protomer restores wild-type pause durations (Fig. 4(A)), as well as wild-type step sizes (Fig. S7). So by removing the ability of the WB protomer to bind flanking DNA, we have restored normal remodeling behavior and eliminated the dominant negative effect of the WB mutation. This suggests that ATP hydrolysis by both protomers promotes their coordinated sensing of flanking DNA and efficient mobilization of the nucleosome.

**Figure 4:**
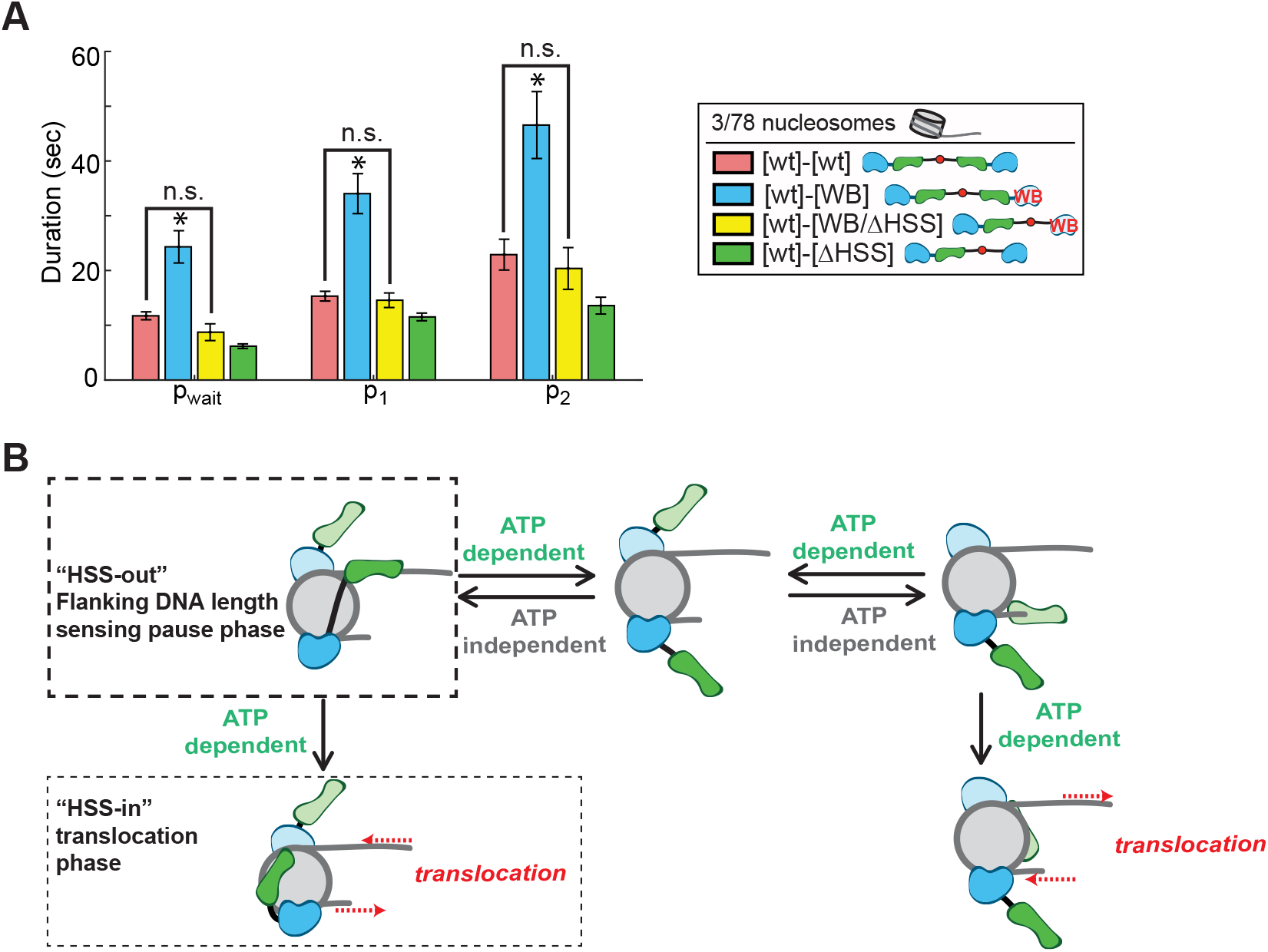
ATP hydrolysis is required for the two motors to coordinate their length sensing activities in the pause phase. (A) Average durations of the first three pauses when [wt]-[wt] or three asymmetric mutants remodel 3/78 nucleosomes, for ∼100 nucleosomes each. All enzyme concentrations are saturating (25 nM [wt]-[wt], 100 nM [ΔHSS]-[wt], 50 nM [wt]-[WB], 200 nM [WB/ΔHSS]-[wt]); ATP is saturating (1 mM). [wt]-[wt] and [wt]-[WB] data are the same as in Fig. 3(A) and are shown here for comparison. Errors were bootstrapped as described in the Methods. (B) Model for how ATP hydrolysis is used to coordinate the flanking DNA length sensing activities of the two SNF2h protomers.

Finally, we also generated a construct in which both protomers are wild-type for ATP hydrolysis, but only one protomer has a DNA-binding domain. This construct, called [wt]-[ΔHSS], actually has slightly shorter pauses than [wt]-[wt], suggesting that when a protomer does not need to coordinate with its partner, it might be even more efficient at pause exit (Fig. 4(A)). All other features of the [wt]-[ΔHSS] construct are wild-type (Fig. S7).

## 3 Discussion

Many chromatin remodeling enzymes that slide nucleosomes are regulated by the length of DNA flanking the nucleosome. Here we show that when a dimeric ISWI family remodeler has the capacity to sense flanking DNA on *both* sides of the nucleosome, ATP hydrolysis by both protomers is required to coordinate this dual DNA length sensing and to gate the nucleosome sliding reaction (Fig. 4(A)).

We previously described a large conformational change in SNF2h as a function of nucleotide state, and had speculated that this conformational change gates pause exit at the single-nucleosome level ([31]). Since the conformational change in SNF2h appears to coincide with ATP hydrolysis, we attributed the ATP dependence of the pause phases observed by smFRET to the ATP hydrolysis needed to promote the conformational change. The single-nucleosome resolution data presented here indicates an additional role for ATP hydrolysis during the pause phase. We therefore postulate that ATP hydrolysis during the pause phase promotes movement of the HSS domain to allow partitioning between two outcomes: (1) the complete release of flanking DNA by the HSS, allowing the other protomer to take its turn, or (2) binding of the HSS domain to the nucleosome core, enabling a translocation-competent state ([31]). The first outcome ensures that neither protomer spends too long with its HSS bound to flanking DNA if it cannot undergo the conformational change to the translocation competent state. Prior work has indicated that under saturating conditions, ATP hydrolysis by ISWI enzymes increases with increasing flanking DNA length ([21, 37, 38]). We therefore further speculate that when the HSS is bound to the longer flanking DNA, ATP hydrolysis is more productive, partitioning a greater proportion of the enzyme to the translocation competent state.

In the model described above (Fig. 4(B)), the dominant negative effect of the WB mutation in one protomer is due to a stymying of outcome 1. When the HSS domain of the catalytically compromised protomer is bound to DNA, the system gets stuck in an unproductive state, in which the wild-type promoter is unable to bind the flanking DNA on its side of the nucleosome, hydrolyze ATP, undergo the conformational change and slide the nucleosome. However, when the catalytically compromised protomer does not have an HSS domain with which to bind flanking DNA, it cannot impede the wild-type protomer, and so the [wt]-[WB/ΔHSS] construct remodels nucleosomes like [wt]-[wt].

In combination with previous studies, our results thus demonstrate that ATP is used not just to physically slide the nucleosome in the translocation phase, but for at least two aspects of the *regulation* of this sliding activity in the pause phases of the reaction ([11, 31, 12]). This use of ATP for regulation is shared with at least one other family of remodelers, INO80 ([15]). It is interesting to speculate about how this ATP requirement for regulation, not just activity, would impact a cell’s control of its chromatin state under limiting ATP conditions (John Tamkun, personal communication).

The directional nucleosome sliding exhibited by ISWI remodelers, regulated not just by flanking DNA but by a comparative assessment of the relative lengths of DNA flanking both sides of a nucleosome, is an important biological function implicated in the generation of evenly spaced nucleosome arrays at TAD boundaries, heterochromatin regions, and in transcriptional regulation ([25, 27, 28, 23, 24, 39, 26]). Not only is this process tightly regulated, but it is also modulatable through the addition of non-catalytic accessory subunits to form complexes with varying properties and, presumably, varying in vivo roles ([40]). For example, the Acf1 accessory subunit forms a complex with SNF2h that is sensitive to longer lengths of flanking DNA than SNF2h alone, but other complexes of SNF2h with different accessory subunits, such as Rsf1 (in the RSF complex), do not have the same DNA length sensitivity ([40]).

Similarly, we show here that the SNF2h motor subunit alone is not processive (Fig. 2), but the ACF complex is ([11]). If the Acf1 accessory subunit modulates the processivity of the motor subunit, it is possible that other accessory subunits that form complexes with SNF2h may modulate its processivity in other ways. While the ability to modulate the length of flanking DNA to which the enzyme is sensitive might change aspects of static structures that are generated (e.g., the precise nucleosome spacing achieved), modulating processivity changes the local *dynamics*. A highly processive enzyme will remain bound at a locus and continually slide the nucleosome back and forth, increasing, for example, the local DNA accessibility in the region to which that complex is targeted. The potential for ATP-dependent chromatin remodelers to mark active chromatin through specific nucleosome dynamics, not just through static chromatin structures, has been recently reviewed in [39]. Interestingly, the synthetic, constitutively dimeric [wt]-[wt] construct recapitulates the increased processivity of the ACF complex, without compromising the ability of the two motors to coordinate their activities. It is possible that the enhanced processivity of both ACF and [wt]-[wt] is due to a decrease in dissociation rate; ACF and [wt]-[wt] both have tighter affinities for nucleosomes than SNF2h alone ([31, 30, 29]).

SNF2h is a complex enzyme, with multiple “moving parts” (the HSS domain discussed here, as well as others [[9, 31]]), that carries out a tightly regulated nucleosome sliding reaction in response to flanking DNA on both sides of the nucleosome. By combining a biochemical system for generating asymmetrically mutant dimers of SNF2h with the ability to watch these dimers remodel single nucleosomes, we can gain mechanistic insights that are difficult to obtain by ensemble enzymological approaches alone. smFRET assays have now been developed for members of all four major remodeler families ([15, 41, 11, 12, 13, 42, 14, 43, 36]), and we anticipate many exciting new avenues of investigation for chromatin remodeling enzymes as this assay becomes part of the standard enzymological toolkit.

## 4 Methods and Materials

### 4.1 Nucleosome labeling and reconstitution

Nucleosomal DNA was generated by PCR as in [15] and [9]. DNA sequences are shown in Fig. S1. HPLC-purified, biotinylated primers were purchased from IDT (Coralville, IA); HPLC-purified, Cyanine5 SE end-labeled primers were purchased from TriLink Biotechnologies (San Diego, CA); and a PAGE-purified primer with an internal Cy5 for generating the 60/60 construct was purchased from IBA Life Sciences (Gottingen, Germany). All nucleosomes were assembled on the 601 nucleosome positioning sequence ([44]).

Recombinant *X. laevis* histones were expressed in and purified from *E. coli* as in [15] and [9]. Histone octamer was assembled as in [15] and [9] with a 2:1 unlabeled:labeled H2A mixture, with the labeled H2A containing a Cy3 attached to an engineered cysteine at position 120 via cysteine-maleimide chemistry. The one endogenous cysteine at position 110 of histone H3 was mutated to an alanine. Nucleosomes were assembled from histone octamer and DNA by salt gradient dialysis over 40-60 hours at 4°C, purified by glycerol gradient centrifugation, and quantified by native gel as in [15].

### 4.2 Remodeler purification

Wild-type SNF2h was purified from *E. coli* as in [9]. The synthetically connected dimeric constructs were generated as in [31].

### 4.3 Native gel remodeling assay

The remodeling reactions quantified by native gel in Fig. S3 were performed at 20°C under single turnover conditions (enzyme in excess of nucleosomes), with 15 nM 60/60 nucleosomes, and saturating enzyme (103 nM SNF2h or 51 nM [wt]-[wt]). Reactions were performed in 70 mM KCl, 1.41 mM MgCl_2_, 0.02% NP-40, 12 mM Hepes-KOH (pH 7.5 at 20°C), 0.1 mM EDTA, and 7% glycerol. Reactions were assembled without ATP and incubated for 5 minutes at 20°C. Reactions were then initiated by the addition of ATPoMg to a final concentration of 1 mM; timepoints were quenched in an equal volume of stop solution (0.6 mg/mL stop plasmid, 40 mM ADP, 16% glycerol). Timepoints were resolved by native PAGE (6% acrylamide, 0.5X TBE) and imaged on a Typhoon variable mode imager by scanning for Cy5 intensity. “% unremodeled” was quantified in ImageJ as the background-corrected intensity of the band that migrated at the same position as the nucleosomes alone lane, divided by the background-corrected intensity of the entire lane.

### 4.4 Ensemble FRET remodeling assay

Ensemble FRET measurements were made on a K2 fluorimeter (ISS, Champaign, IL) by following the intensity of Cy5 over time (see also [15, 9]). Excitation and emission wavelengths were 515 nm and 670 nm respectively, with a 550 nm short pass filter in the excitation path and a 535 nm long pass filter in the emission path. Emission intensities were measured every second.

All reactions were performed under single-turnover conditions with saturating enzyme and saturating (1 mM) ATP at 20°C. Nucleosome concentrations were 7.5 nM; enzyme concentrations are given in figure legends. Reaction conditions were 12 mM HEPES-KOH, pH 7.5 at 22°, 70 mM KCl, 1.41 mM MgCl_2_, 7% glycerol, 0.02% Igepal (Spectrum Chemical I11112, New Brunswick, NJ), 0.1 mM EDTA, and 1 mM ATP-Mg. Remodeling was initiated by the addition of enzyme and ATP.

Cy5 intensities were normalized by fitting the unnormalized intensities to a two-phase exponential decay with 5 free parameters. In particular, the intensity of Cy5 at time *t, I*(*t*), was fit to

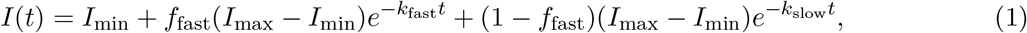

where *I*_min_ and *I*_max_ are the minimum and maximum intensities respectively, *k*_fast_ and *k*_slow_ are the rate constants of the fast and slow phases of the reaction respectively, and *f*_fast_ is the fraction of the reaction in the fast phase. Cy5 intensities were then normalized to the fitted *I*_min_ and *I*_max_ values. The same fitting and normalization were performed on the “pseudoensemble” data in Fig. S6.

### 4.5 Single molecule FRET assay

smFRET measurements were performed as in [15], except that the SNF2h-specific buffers in [9] were used instead of the INO80 buffers in [15]. The final remodeling reaction conditions (Imaging Buffer) were 53 mM HEPES-KOH, pH 7.5 at 22°C, 9.1 mM Tris-acetate, pH 7.5 at 22°C, 63 mM KCl, 1.4 mM MgCl_2_, 10% glycerol, 0.1 mM EDTA, 0.02% Igepal, 1% glucose, 0.1 mg/mL acetylated BSA (Promega, Madison, WI), 2 mM Trolox (Sigma 238813), 0.03 mM *β*-mercaptoethanol, 2 U/*μ*L catalase (Sigma E3289), and 0.08 U/*μ*L glucose oxidase (Sigma G2133). Remodeling was initiated by the addition of ATP and enzyme at the concentrations indicated in figure legends using an automated syringe pump (J-KEM Scientific, St. Louis, MO). Images were collected using Micro-Manager (www.micro-manager.org, San Francisco, CA) ([45]) at 7.4 Hz, with an exposure time of 100 ms. For the chase experiments for measuring processivity in Fig. 2, the chamber was flushed with a large volume (600 *μ*L) of Imaging buffer with 1 mM ATP but no enzyme, again using the automated syringe pump, 50 s after the first injection that initiates remodeling (see also [15]).

FRET-versus-time trajectories were extracted from microscope images using our custom Matlab software package, Traces (https://github.com/stephlj/Traces) ([15, 46]). Pause durations were quantified using a hidden semi-Markov model as in [15], through the adaptation of the pyhsmm python library (https://github.com/mattjj/pyhsmm) that accompanies the Traces software package.

As in [15] and [9], only nucleosomes that started in the higher-FRET, proximally-labeled cluster (see Fig. S2), and only those that exhibited single-step photobleaching of both dyes, were retained for further analysis. Except for data used to quantify processivity, only the portions of trajectories prior to any direction reversals and prior to the nucleosome being moved out of FRET range (defined as 0.275 FRET) were retained for further analysis (quantification of pause durations, step size, etc).

All errors were estimated by bootstrapping over FRET-versus-time trajectories. For pause durations, this means sampling all the trajectories in a dataset with replacement 1000 times, recomputing the average pause duration with this new sampled dataset, and then taking the standard deviation of these 1000 new means as the error. A similar process was used to estimate the error on step size CDFs.

#### Quantifying enzyme processivity

To quantify enzyme processivity in Figs. 2 and S4, we counted the number of transitions between pause states as a function of time. A processive enzyme will continue to transition between pause states (that is, translocate the nucleosome to different positions along the DNA) even under chase conditions.

We count the total number of nucleosome translocations in a trajectory as the number of pause states minus one. We first quantified pauses by a hidden Markov model (HMM) using the pyhsmm package, as described above. We then hand-curated each trajectory to remove obviously spurious state transitions. This hand curation was necessary because these processivity experiments required measuring very long trajectories, which get assigned more noise states by the HMM (the longer a trajectory, the more likely it will have noise events to which the HMM incorrectly assigns real states). Also, the position of the Cy5 dye at 9 bp outside of the nucleosome, rather than 3 bp, and the continual remodeling of the nucleosome in and out of FRET range, means that significant portions of these traces are at low FRET values, which tend to be inherently more noisy. The noise states that were removed were often of very short duration (several frames) and assigned nonsensically high or low FRET values by the HMM, and so were clearly spurious.

The cumulative number of nucleosome translocations was counted in 20-second intervals for each trajectory. Errors on the average cumulative translocations were estimated by a bootstrapping approach: trajectories in a dataset were resampled with replacement 1000 times, the cumulative number of nucleosome translocations at 20-second intervals was counted for each resampled dataset, and the standard deviation taken as the error.

#### FRET-to-bp calibration

The calibration that converts FRET values to base pairs of DNA that the nucleosome has been slid were performed as in [15], except that the Cy3 donor dye was on histone H2A at position 120 rather than on histone H3 at position 33. This different histone labeling position resulted in a different relationship between FRET and flanking DNA length (Fig. S2(A)). In fact, the relationship between FRET and *n*, the number of bp between the Cy5 label and the edge of the nucleosome, was so different that it was not well described by the expression used in [15] to fit the calibration curve data in that work.

In [15], we found that the best expression to relate FRET and *n* was

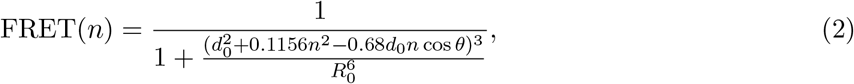

where *R*_0_ is the Forster radius for Cy3-Cy5 in nm ([48, 47]) and *d*_0_, *n* and *θ* are defined in Fig. S2(B). In [15], we also considered two variations on Eq. 2. First, because zero FRET is difficult to measure, we considered a constant offset, FRET_0_, which gave the relationship

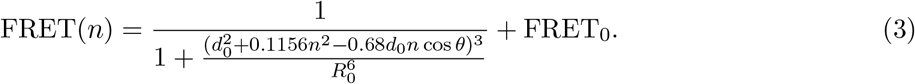

However, because this significantly increases the number of free parameters, we reduced the dimensionality of the fit to one comparable to Eq. 2 by assuming that, given the geometry suggested by the crystal structure of the nucleosome, *θ* ≈ 90°. In that case,

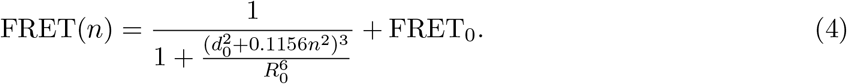

In Fig. S2(C), we show that Eqs. 3 and 4 better describe the relationship between FRET and *n* for the H2A-labeled nucleosomes used in this work. Fit parameters are given in Table 1; note that as in [15], we performed a global fit to the data for the proximal and distal FRET clusters, which have separate *d*_0_, *θ*, and FRET_0_ parameters, but share the same *R*_0_. Errors on fit parameters were obtained by a bootstrapping routine as in [15].

**Table 1:** Parameters obtained by fitting Eqs. 2, 3, or 4 to the calibration data in Fig. S2. Eq. 4 is the one used in this work. The “prox” and “dist” subscripts refer to the proximal and distal peaks in the FRET KDEs in Fig. S2(A). Errors are bootstrapped as described in the Methods. ^1^[47].

| Fit to: | $R_0$ (nm) | $d_{0,\text{prox}}$ (nm) | $\theta_{\text{prox}}$ ($^\circ$) | $\text{FRET}_{0,\text{prox}}$ | $d_{0,\text{dist}}$ (nm) | $\theta_{\text{dist}}$ ($^\circ$) | $\text{FRET}_{0,\text{dist}}$ |
| --- | --- | --- | --- | --- | --- | --- | --- |
| (expected) | $6^1$ | $\sim 2$ | $> 90$ | | $\sim 6$ | $\sim 90$ | |
| Eq. 2 | $13 \pm 1$ | $8 \pm 1$ | $180 \pm 12$ | (N/A) | $11 \pm 1$ | $120 \pm 45$ | N/A |
| Eq. 3 | $7 \pm 2$ | $5.3 \pm 0.8$ | $110 \pm 26$ | $0.14 \pm 0.03$ | $6.8 \pm 0.9$ | $87 \pm 8$ | $0.07 \pm 0.08$ |
| Eq. 4 | $5.7 \pm 0.3$ | $4.6 \pm 0.2$ | (90) | $0.16 \pm 0.03$ | $5.6 \pm 0.2$ | (90) | $0.17 \pm 0.02$ |

Many of the fit parameters for Eqs. 3 and 4 are close to expected values and within error of each other. In particular, both *θ*_prox_ and *θ*_dist_ are within error of 90°, which would suggest Eq. 4 is the more reasonable expression. Finally, both expressions have essentially identical behaviors for the proximal FRET data between 3 and 25 bp, the range of values we care most about. Therefore we use Eq. 4 to convert between FRET and *n* in this work.

As in [15], we invert Eq. 4 to obtain

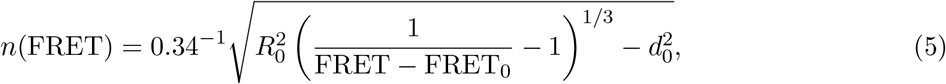

which we use to convert from measured FRET value to bp the nucleosome has been moved.

## 5 Acknowledgments

We thank Julia Tretyakova for histone purifications, the Narlikar lab for helpful discussions, Nathan Gamarra for advice on experimental design and on this manuscript, and Matthew Johnson for help with statistical analyses and adapting the pyhsmm package for our system. This work was supported by NIH grants to G.J.N. (R01GM073767 and R35 GM127020) and a Leukemia and Lymphoma Society Career Development Program Fellow award to S.L.J.

## 6 Supplemental Figures

**Figure S1:**
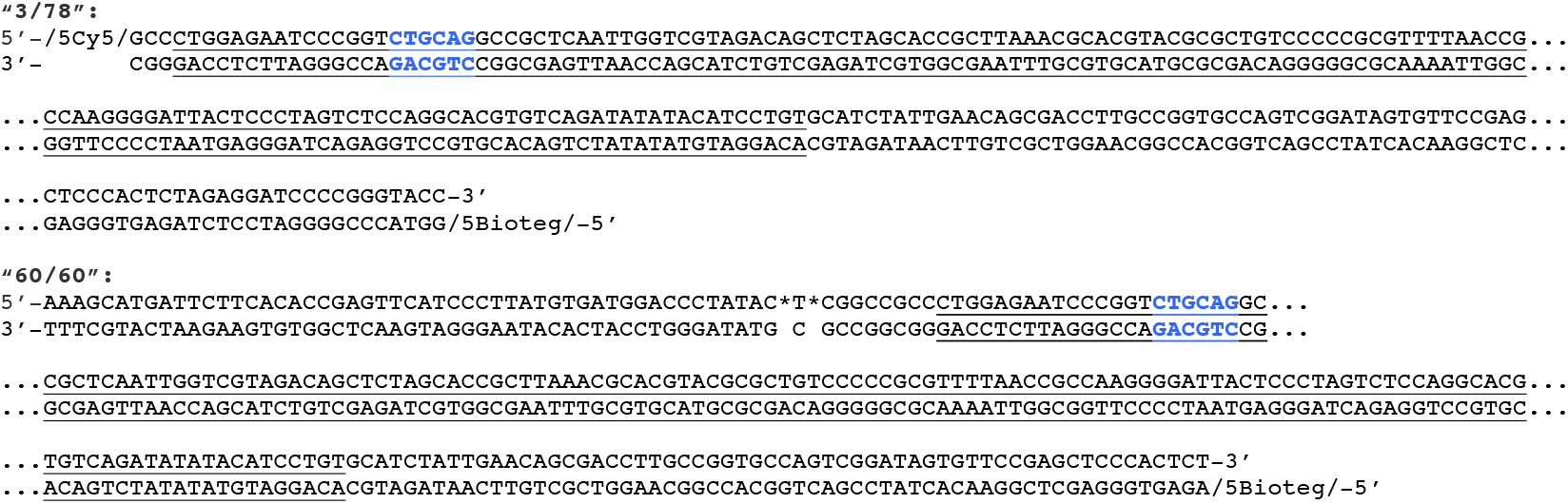
Supplement to Fig. 1: DNA sequences used in this work. Sequences of nucleosomes that start end-positioned (“3/78”, top) or centered (“60/60”, bottom). The 601 positioning sequence ([44]) is underlined. A Pst1 restriction site has been engineered into the 601 sequence 18 bp from one end, shown in bold blue letters. The base with an internal Cy5 label in the 60/60 construct is flanked by asterisks. DNAs used for the calibration curve (Fig. S2) were generated by adding bp to the Cy5-labeled short end of the 3/78 construct, using the same sequence as in the 60/60 construct; see also [15].

**Figure S2:**
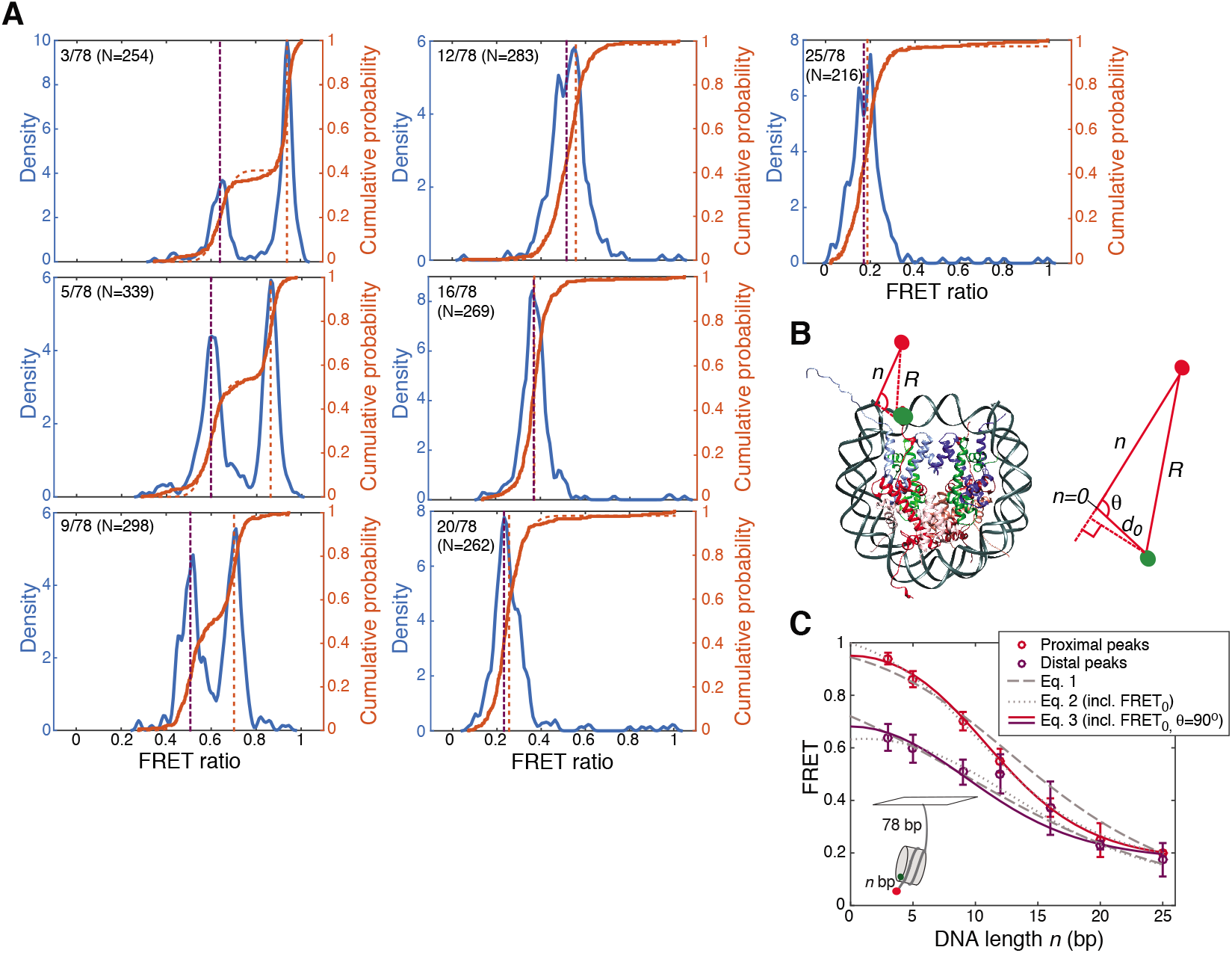
Supplement to Fig. 1: (A) FRET values of the nucleosome constructs used to generate calibration data for converting FRET to bp of DNA moved out of the nucleosome. Constructs are labeled as *n*/78, with 3≤ *n* ≤ 25 bp on one side of the nucleosome, and 78 bp on the other (see schematic in (C)). N is the number of nucleosomes in each data set. Blue curves are KDEs with bandwidth 0.01; solid orange curves are corresponding empirical CDFs; dashed orange curves are fits to a Gaussian mixture model for identifying peak positions and widths. FRET values for nucleosomes cluster into two populations, a “proximal” population, in which Cy3 is on the H2A closest to the Cy5 label (as in (B)), and a “distal” population, in which the other H2A is labeled, resulting in a lower FRET value because the dyes are further apart. Nucleosomes with neither or both H2A’s labeled are excluded. See [15] for more details. (B) Parameters used in derived calibration curve models. *n* is the number of bp of DNA between the edge of the nucleosome and the Cy5 label, *R* is the distance between the Cy3 and Cy5 dyes in three dimensions, *d*_0_ is the distance between the dyes when *n* = 0, and *θ* is the angle between *d*_0_ and the flanking DNA. (C) Fits of Eqs. 2, 3, and 4 to FRET values of proximally and distally labeled nucleosomes, as a function of *n*. Fit parameters are given in Table 1. Errors on the data are standard deviations from fits of Gaussian mixture models to the CDFs of FRET values for each nucleosome construct in (A).

**Figure S3:**
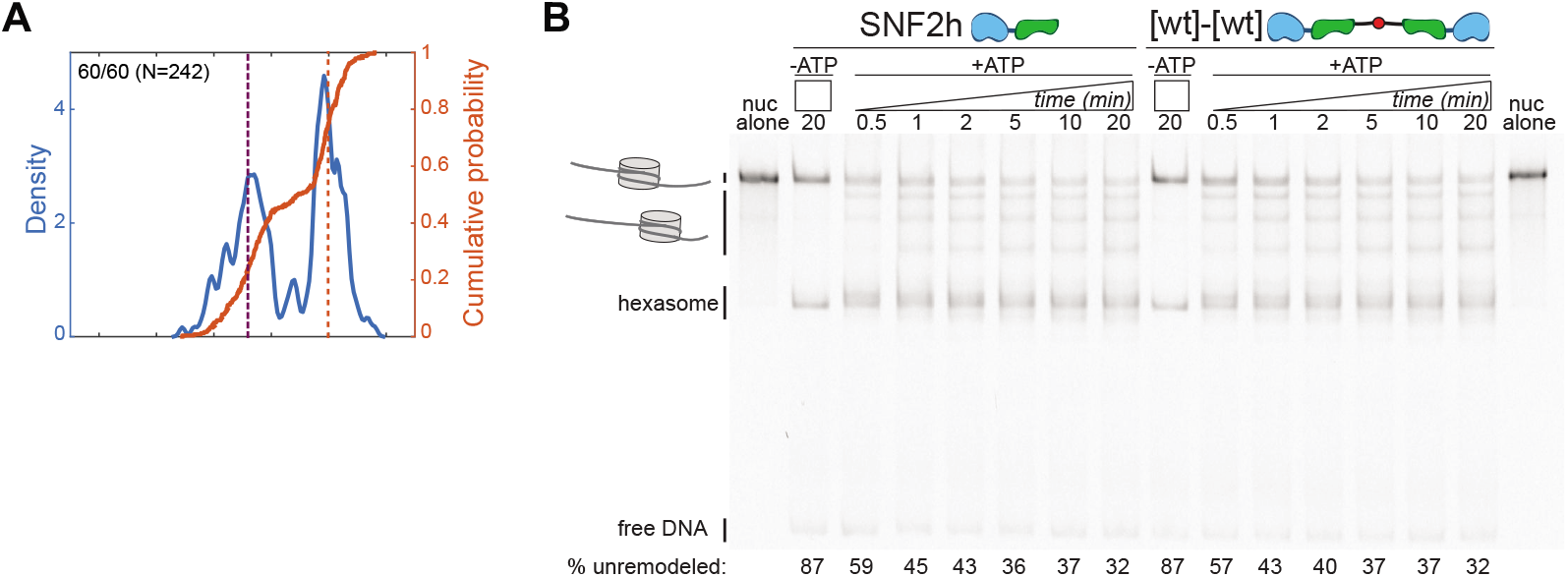
Supplement to Fig. 2: (A) KDE and CDF of starting FRET values for the 60/60 nucleosomal construct (see Fig. S2(A)). (B) Gel remodeling of 60/60 nucleosomes by 103 nM SNF2h or 51 nM [wt]-[wt], with 1 mM ATP. After 5 minutes, ~40% of the nucleosome population is still centered (the smFRET reaction is maximally ~6 min, due to photobleaching). Note that SNF2h and [wt]-[wt] have the same product distributions on 60/60, indicating that although [wt]-[wt] might be more processive on these nucleosomes like ACF, it does not have the enhanced length sensitivity of ACF (ACF would keep these nucleosomes centered ([29]).

**Figure S4:**
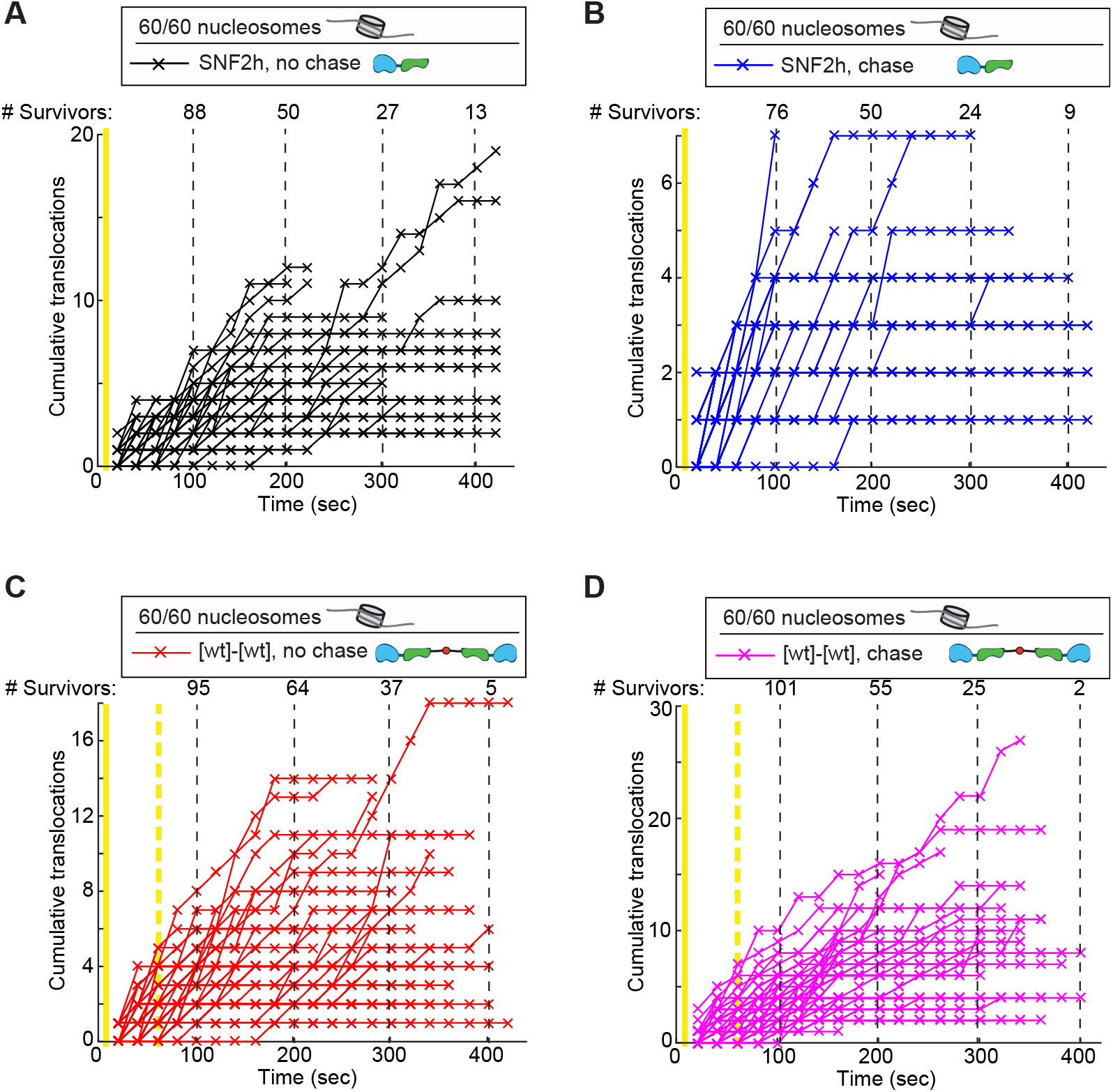
Supplement to Fig. 2: Cumulative translocations in individual trajectories for (A) SNF2h, no chase, (B) SNF2h, with chase, (C) [wt]-[wt], no chase, and (D) [wt]-[wt], with chase. Each panel has a different y-axis scale. “# Survivors”: Number of trajectories at each indicated time point that have not photobleached; for example, the SNF2h, no chase sample starts with 103 trajectories, but after 400 seconds, all but 13 trajectories have photobleached. See Fig. 2 for conditions.

**Figure S5:**
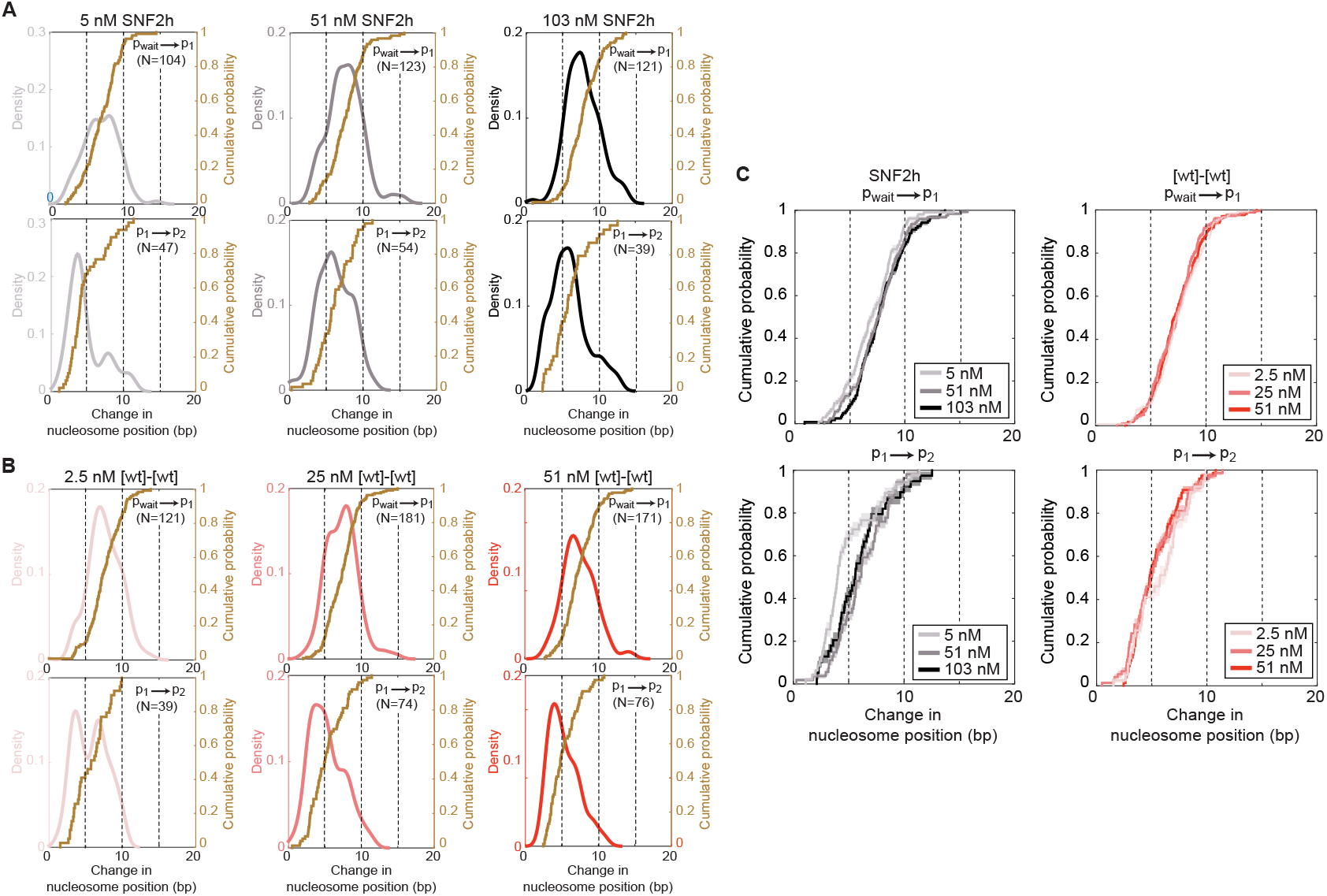
Supplement to Fig. 2: Translocation step sizes of SNF2h and [wt]-[wt] at saturating versus subsaturating enzyme concentrations (and 1 mM ATP). (A,B) KDEs of translocation step sizes, with corresponding CDFs overlaid, for varying concentrations of SNF2h (A) or [wt]-[wt] (B). See Fig. 1(D) for a description of these plots. N: number of remodeling events included in the KDE and CDF. KDE bandwidths are 0.75. (C) Overlaid CDFs for the first (top) or second (bottom) translocation events, at the varying concentrations in (A). Mean step sizes for first and second translocations respectively are: 7.0±0.2 bp and 4.8±0.4 bp for 5 nM SNF2h; 7.6±0.2 bp and 6.2±0.3 bp for 51 nM SNF2h; 7.8±0.2 bp and 5.8±0.4 bp for 103 nM SNF2h; 7.5±0.2 bp and 5.7±0.4 bp for 2.5 nM [wt]-[wt]; 7.3±0.2 bp and 5.3±0.3 bp for 25 nM [wt]-[wt]; 7.4±0.2 and 5.3±0.2 bp for 51 nM [wt]-[wt].

**Figure S6:**
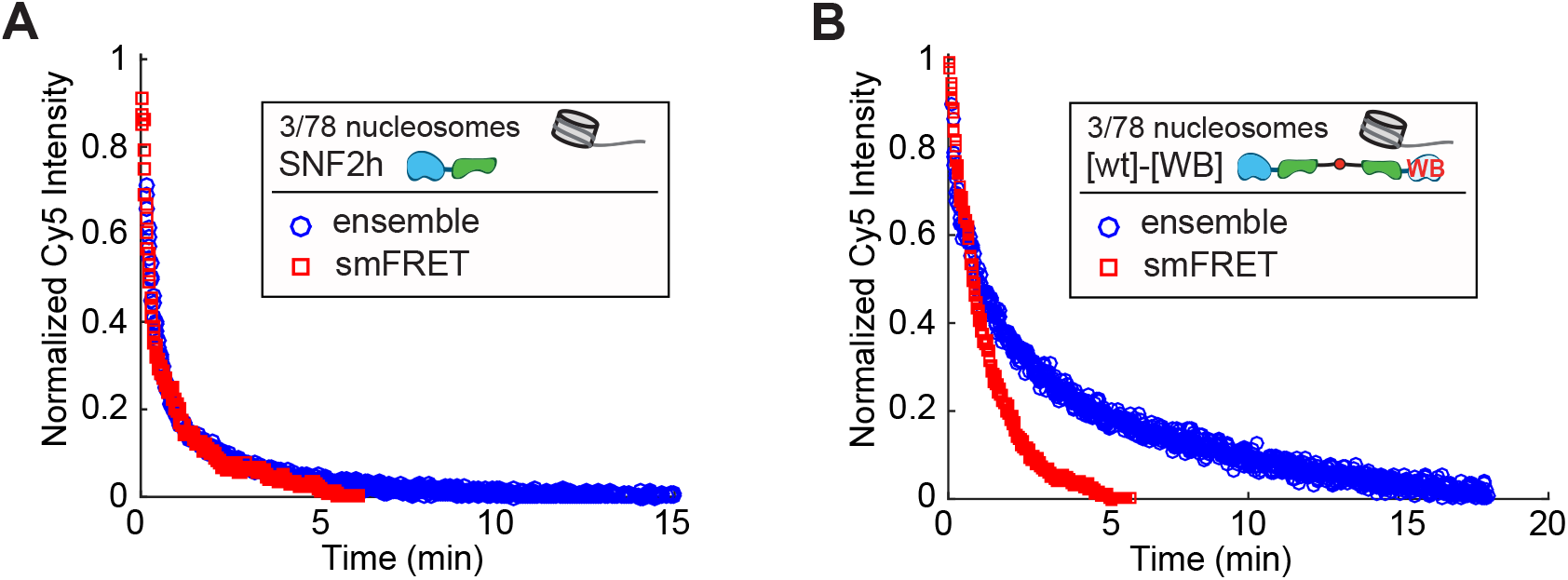
Supplement to Fig. 4: (A) Ensemble FRET remodeling data (measured as the decrease in overall sample Cy5 intensity as a function of time; blue data), compared to a “pseudoensemble” measure of the overall remodeling rate by single-molecule FRET (red data), for 103 nM SNF2h plus 1 mM ATP. The “pseudoensemble” data are the summed Cy5 intensities of all surface-attached nucleosomes in the indicated single molecule data set, binned in 1-second intervals to simulate the measurement of ensemble Cy5 intensities. Consistent with previous work, the overall remodeling rates measured by smFRET and by ensemble FRET are comparable ([11, 13]). (B) Same as (A) but for 50 nM of the [wt]-[WB] construct. Here, the overall reaction rates measured by ensemble FRET versus smFRET are not the same; in particular, the smFRET rate appears faster than the ensemble rate. As in our previous work with acidic patch mutant nucleosomes ([9]), this discrepancy is due to photobleaching in the smFRET reaction, which masks slowly remodeled nucleosomes. Thus the pause durations reported for the [wt]-[WB] construct in Figs. 3 and 4 are a lower bound on the actual pause durations.

**Figure S7:**
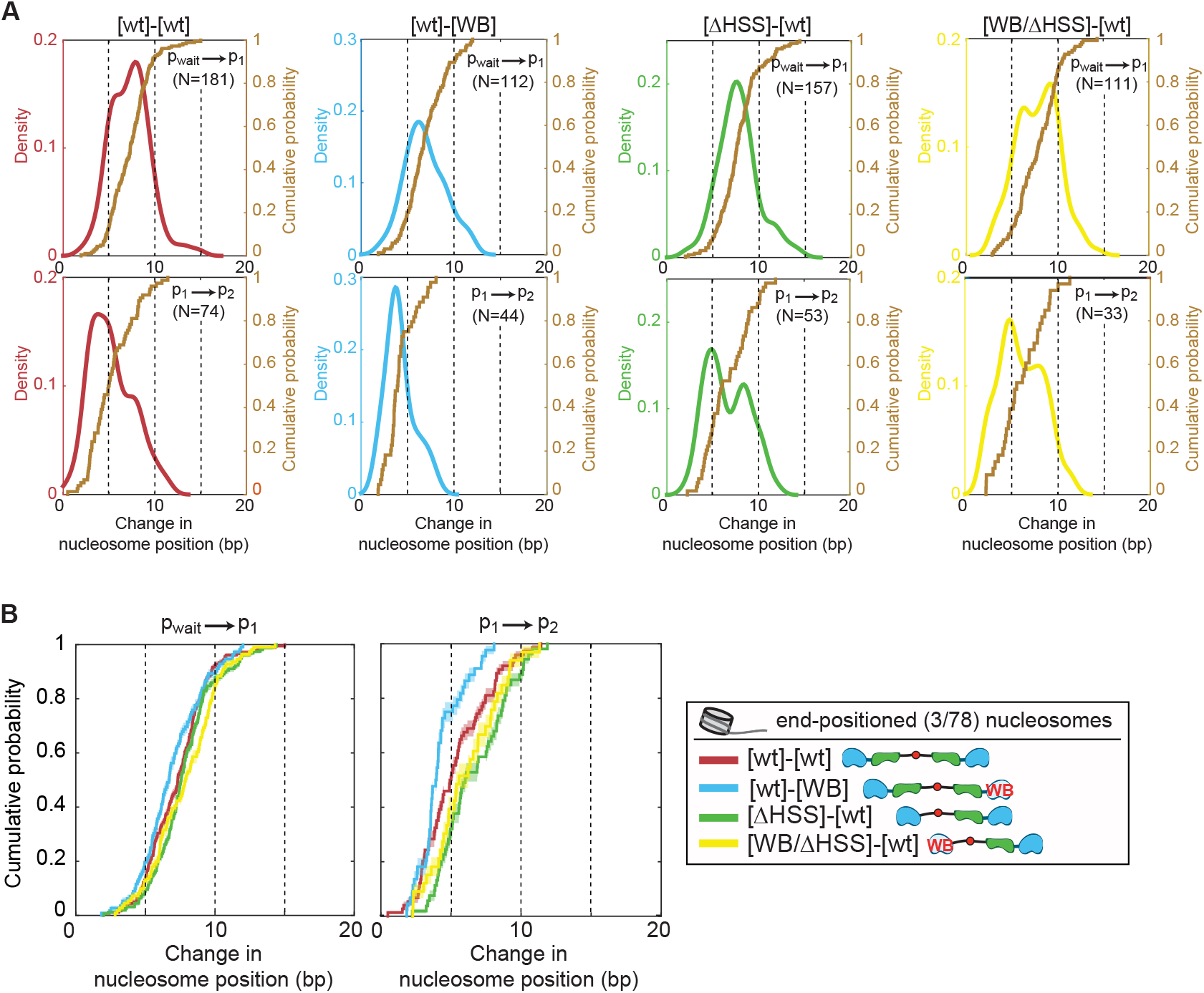
Supplement to Fig. 4: Translocation step sizes of mutants. (A) KDEs of translocation step sizes, with corresponding CDFs overlaid, for the [wt]-[wt] construct and the four asymmetrically mutant dimers. See Fig. 1(D) for a description of these plots. N: number of remodeling events included in the KDE and CDF. KDE bandwidths are 0.75. (B) Overlaid CDFs for the first (left) or second (right) translocation events, for the four constructs in (A). Mean step sizes for first and second translocations respectively are: 7.3±0.2 bp and 5.3±0.3 bp for [wt]-[wt]; 6.9±0.2 bp and 4.2±0.2 bp for [wt]-[WB]; 7.7±0.2 bp and 6.6±0.3 bp for [ΔHSS]-[wt]; 7.8±0.2 bp and 6.1±0.4 bp for [WB/ΔHSS]-[wt]. All enzyme concentrations are saturating (25 nM [wt]-[wt], 100 nM [ΔHSS]-[wt], 50 nM [wt]-[WB], 200 nM [WB/ΔHSS]-[wt]); ATP is saturating (1 mM).

